# Human photoreceptor cell transplants integrate into human retina organoids

**DOI:** 10.1101/2022.08.09.500037

**Authors:** Felix Wagner, Roberto Carrera, Thomas Kurth, Stylianos Michalakis, Ronald Naumann, Marta Zuzic, Katrin Neumann, Olivier Gourau, Volker Busskamp, Mike O. Karl

## Abstract

Cell transplantation is a promising therapeutic approach to recover loss of neurons and vision in patient retinas. So far, human photoreceptor transplants restored some visual function in degenerating mouse retina. Whether retinal cell transplants also integrate into human retina, and how to optimize this for different pathologies are still unknown. Here, we sought to determine if human retina organoids generated from pluripotent stem cells might assist cell replacement therapy development in a human-to-human setting. Models for intra- and subretinal cell transplantation strategies were explored: Photoreceptor donor cells carrying a transgenic fluorescent reporter were enriched from acutely dissociated human retinal organoids. Donor cells were precisely transplanted by microinjection into the retina of host organoids, but high cell numbers might require multiple injections posing potential damage. Alternatively, donor cells were transplanted in large numbers by placing them in subretinal-like contact to the apical organoid surface. Using postmitotic retinal organoids (age >170-days) as a source for donor cells and as hosts, we show that six weeks after subretinal-like transplantation, large clusters of photoreceptors reproducibly incorporate into the host retina. Transplanted clusters frequently are located within or across the host photoreceptor layer, include cone and rod photoreceptors, and become infiltrated by cell processes of host Müller glia, indicative of structural integration. Histological and ultrastructural data of virally-labeled photoreceptor transplants show characteristic morphological and structural features of polarized photoreceptors: inner segments and ribbon synapses, and donor-host cell contacts develop contributing to the retinal outer limiting membrane. These results demonstrate that human retinal organoids provide a preclinical research system for cell replacement therapies.

**Graphical abstract:** 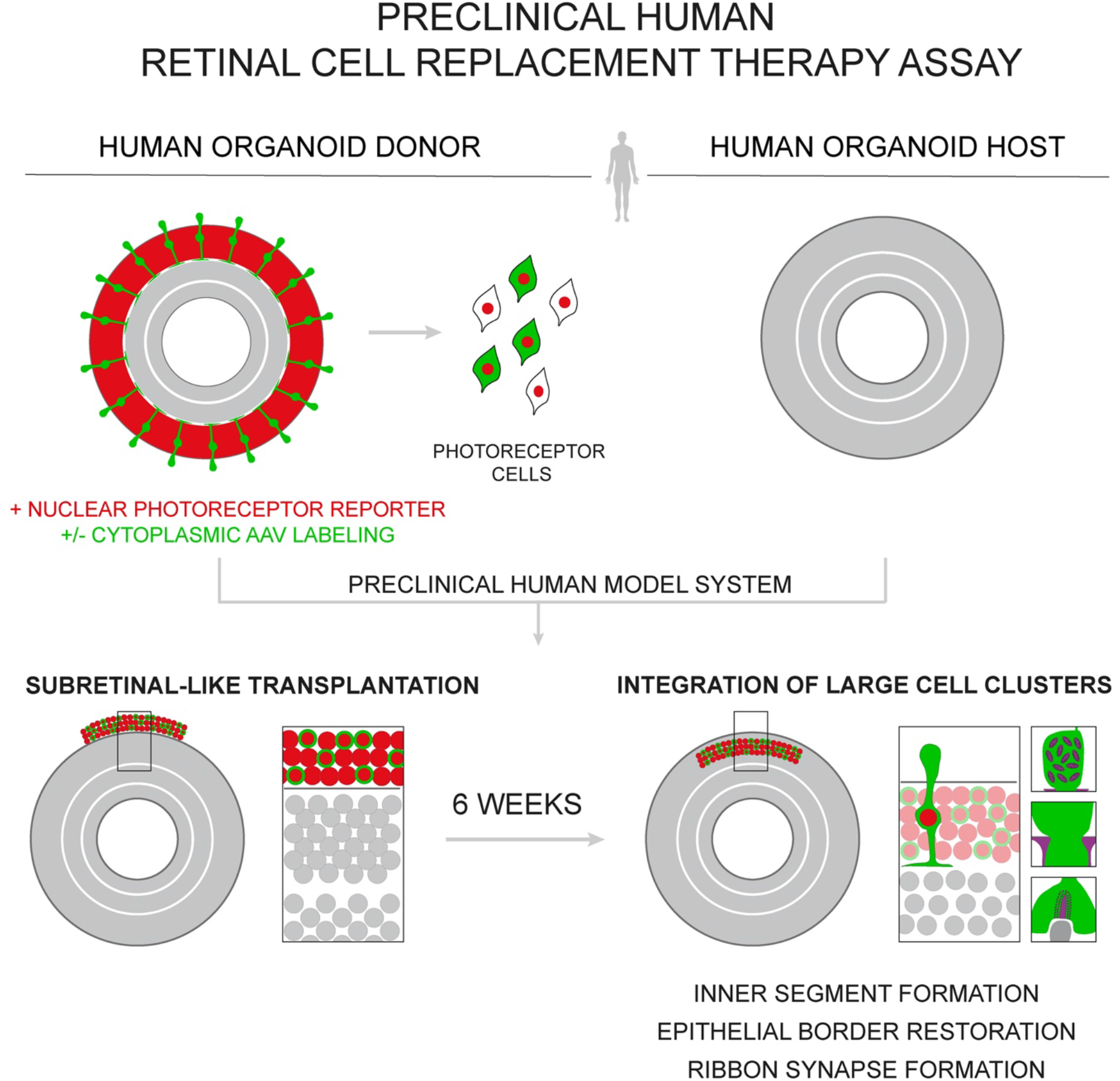

## Introduction

Vision loss is a large personal and socio-economic burden. 61 million people are predicted to be blind worldwide by 2050. Another 474 million people will be affected by moderate to severe vision impairment (1). The ultimate reason for untreatable vision loss is degeneration of the light-sensitive photoreceptor cells within the retina, either as a primary or secondary disease process. Hence, development of therapeutic approaches is crucial to fight neurodegenerative diseases. Gene and drug therapies are currently in development and some already have been approved for specific diseases (2). Cell replacement therapy by transplantation of retinal cells is a promising approach to restore cellular loss and improve visual function. After the very first retinal transplantation studies in the 1980s, the first transplantation of human stem cell-derived photoreceptors was conducted in 2009 (3–5). Former studies could optimize survival and integration of transplanted photoreceptor cells (6, 7). Manipulation of the host retina by disruption of the apical epithelial border of the retina led to a higher number of integrated photoreceptor cells (8, 9). Besides restoration of photoreceptor cell loss, transplantation of rod photoreceptors might in addition protect and thus prevent cone photoreceptor degeneration (10). Recent studies have shown species-specific differences when either mouse or human postmitotic photoreceptor cells are transplanted into mice. Whereas mouse photoreceptors integrate in smaller numbers, if at all, into the host retina (11–13), human photoreceptors, particularly cones, have recently been shown to structurally integrate in large clusters and/or contact host second-order neurons and Müller glia (14, 15). While exchange of genetic and protein material has been shown to frequently occur in mouse-to-mouse photoreceptor transplants (11–13, 16–18), this phenomenon was not observed in human transplants into rodent models (14, 19–21). Hence, animal studies support the possibility of beneficial cell replacement therapy, posing the questions how to effectively translate findings to the clinic, and which patients at what stage of pathology would profit most. Studies in large animal and non-human primates might be key but are limited for larger scale experiments. Nevertheless, non-human primate studies showed the feasibility of transplanting human stem cell-derived retinal cells with indications for cell integration (22, 23). In vivo live imaging approaches even allowed tracking of transplanted cells at single cell resolution (24). Three patient case studies explored retinal transplantation in, and reported graft survival but lack of graft integration up to 3 years after transplantation (25, 26). Together, current data indicate that further optimization of cell transplant products and understanding of the mechanisms regulating and limiting cell integration will be key for clinical translation

Human model systems will be crucial to facilitate clinical translation of cell therapeutic approaches. Organoid technology allows to work in a preclinical context with human models. In recent years, human stem cell derived retinal organoids have been shown to recapitulate development, epithelial structure, and cell types of the human in vivo retina (27–31). Light-induced cell responses and metabolic activity can be measured in organoids by electrophysiological recordings and live imaging (29, 32–34). Whether they can achieve a maturation state where they fully recapitulate photoreceptor function and synaptic wiring is still unclear. Genetic organoid pathology models have been developed from patient-derived stem cells (35–37) whereas pharmacological-induced retinal degeneration has been rarely studied until now (38). Further, gene therapeutic approaches have been established and evaluated in human organoids for their efficient transduction of retinal cells and rescue of gene function (36, 37, 39). Hence human organoid models are already used to model retinal pathologies and to develop gene therapies.

A preclinical human retina system might facilitate cell replacement therapy development and translation. For retinal cell replacement therapy in vivo, transplanted cells can be placed either at the apical site of the retina next to the photoreceptor layer (subretinal), within the epithelial layers of the retina (intraretinal), or at the basolateral site of the retinal epithelium, next to the ganglion cell layer (intravitreal). Until now, there are few studies that investigated whether in vivo transplantation approaches of retinal cells could be experimentally reproduced ex vivo (40, 41), specifically, no studies in human retina ex vivo. To address these questions, we aimed to develop a in vivo-like cell transplantation system in human retinal organoids ex vivo to facilitate development of cell therapeutic approaches in a preclinical human-to-human setting. Here, we aimed to investigate whether transplanted human photoreceptor cells can integrate into the healthy human retina using organoid models.

## Results

### Intra- and subretinal-like cell transplantation models in human retina organoids

To establish a human experimental ex vivo system for cell transplant therapy development, human induced pluripotent stem cell (hiPSC) derived human retinal organoids (HROs) were used both as a source for a transplantable population of human photoreceptor (PR) cells and as a 3D human retina host model (Fig.1A). Using previously established protocols and human pluripotent stem cell lines, we generated HROs (14, 21, 39). Cone PRs derived from such HROs at day 200 were recently shown to be most suitable for transplantation studies in mice (14). 200-days old HROs are postmitotic (14, 39) and recapitulate key features of the human retina in vivo, including several nuclear layers and an apical retinal epithelial boundary, called the outer limiting membrane (OLM) (Suppl. Fig.S1). PR cells have their nuclei in the outer nuclear layer, interneurons and Müller glia (MG) nuclei are found in the inner retina layer (Suppl. Fig.S1). Both layers are each separated by a layer containing synaptic connections (Suppl. Fig.S1). The OLM is formed by cell-to-cell contacts between PRs and MG, and can be visualized by actin-binding phalloidin, with antibodies against zonula occludens protein 1 (ZO1), and against cytoplasmic markers for MG (RLBP1, SCL1A3) as a proxy (Suppl. Fig.S1). To track cell transplants, HROs carrying a transgenic fluorescent PR-specific CRX-mCHERRY reporter (21) were used to acutely isolate single PRs as a suspension by HRO dissociation and fluorescence-activated cell sorting (FACS) (Fig.1A-B; Suppl. Fig.S2A). HROs without fluorescent reporter were used as hosts.

**Fig.1.**
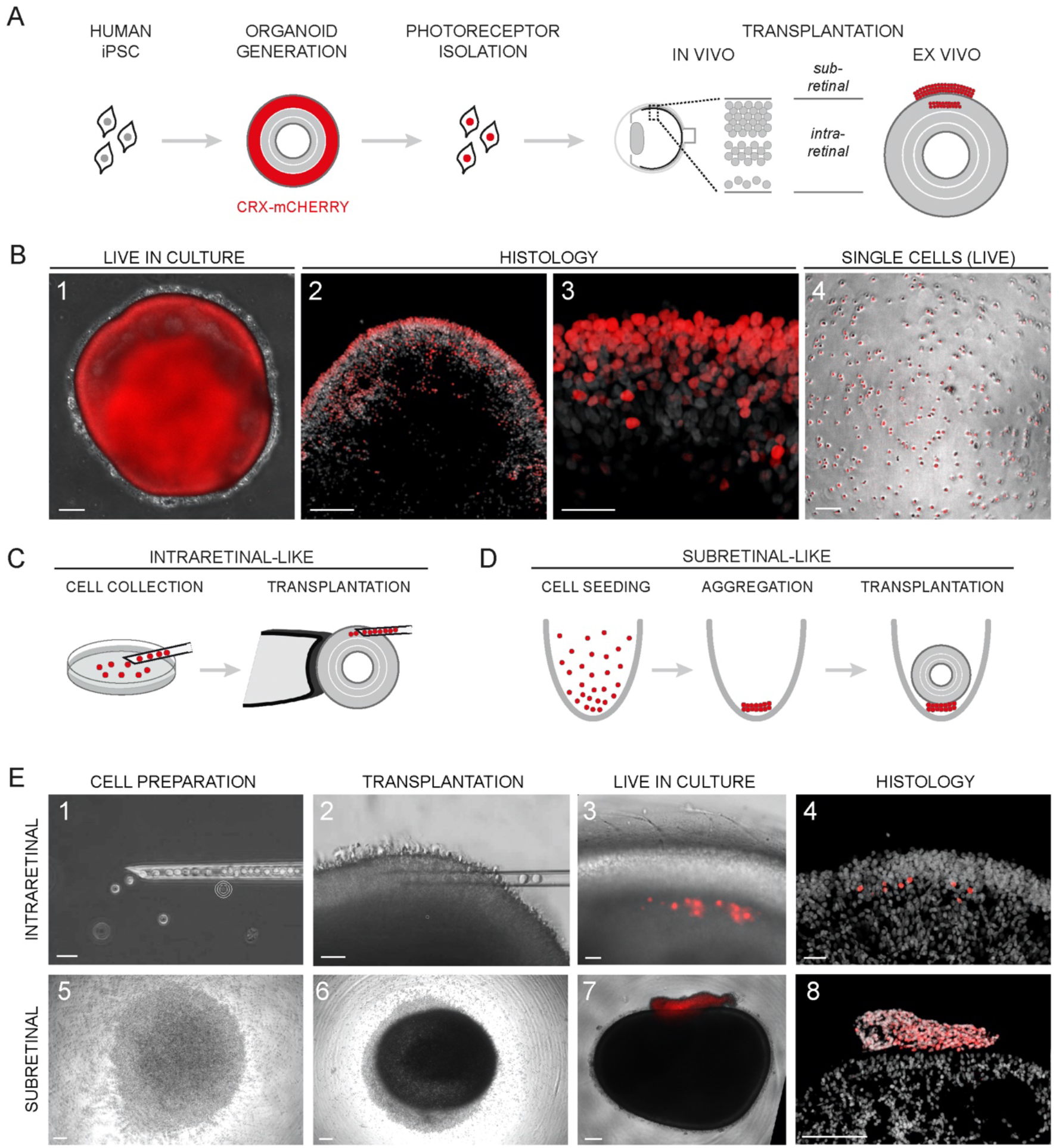
Intra- and subretinal-like cell transplantation models in human retina organoids. (A) Human stem cell-derived retinal organoids (HROs) carrying a transgenic fluorescent photoreceptor-specific CRX-mCHERRY reporter (red) were source for a transplantable cell population. In vivo transplantation approaches into the subretinal space or into the retinal epithelium correspond to ex vivo transplantation onto the HRO surface or injection into the HRO epithelium, respectively. (B) 1: Donor HROs express the photoreceptor-specific CRX-mCHERRY reporter in culture. 2-3: In histology, photoreceptor cells are present in the outer retinal epithelium (DAPI, gray: cell nuclei). 4: The reporter is visible in single cells after HRO dissociation. (C) Intraretinal- and (D) subretinal-like transplantation paradigm using human organoid models. (E) Intraretinal- (1-4) and subretinal-like (5-8) cell preparation and transplantation approaches mimicked ex vivo using HRO hosts. Transplanted cells are visible live in culture (3, 7) and in histological analysis (4, 8) (gray: DAPI, nuclei). Scale bars 100*μ*m (B: 1-2, 4; E: 5-8), 25*μ*m (B: 3; E: 1-4).

As a first approach to precisely position and monitor single PR transplants, we explored utilization of cell microinjection into the retinal epithelial layers of intact living HROs at a postmitotic stage (>day 170) (39) in culture (Fig.1C, E1-2; Suppl. Movies1,2). CRX-mCHERRY labeled transplants were visible directly after injection and upon further HRO culture survived at least for another 3 weeks (Fig.1E3). Histological analysis of HRO sections confirmed this (Fig.1E4). This intraretinal transplantation approach allowed controlled positioning of individual transplanted cells within the host. However, transplantation into several HROs is time- and work-intensive, and to achieve a high number of transplanted cells will require multiple injections posing potential damage to host retina.

Next, we developed an experimental model to target the so far commonly used transplantation site in animal in vivo studies: the subretinal space (Fig.1A, D). To reproduce a subretinal-like contact between transplant and host, i.e. at the apical retinal surface of a host retina, 35,000 HRO-derived single PR cells were injected (pipetted) into round-bottom wells of a 96-well low attachment cell culture plate, which formed a pellet after 2-3 hours (Fig.1E5), followed by placement of a host HRO (>day 170) on top of each transplant (Fig.1D). Live imaging in culture showed that transplant and host HRO come in direct contact (Fig.1E4) as reported for in vivo animal studies (14). After a few days, transplant and host were fixed together and analyzed by histology. Transplanted cells survived, appeared aggregated, and in opposition to the apical border of the host HROs (Fig.1E7-8). However, transplanted cells were clearly still separated from, and did not invade or become interspersed with the host HRO tissue. In summary, we established two methods that potentially reproduce to some extend intra- and subretinal-like PR transplantation in a human retina ex vivo model. This system might be used to evaluate methods to assess and stimulate transplant incorporation, structural and functional integration, and visual function of transplants in healthy and restoration thereof in pathologic host retinas.

### Photoreceptors effectively incorporate after subretinal-like transplantation into HROs

Recent studies in mice in vivo showed that human PRs incorporate in large cell clusters when transplanted into the subretinal space (14), whereas mouse PRs rarely integrate (11–13, 17). To determine whether human PRs incorporate upon subretinal-like conditions into healthy control host HROs (Fig.2A), sections of experimental HROs were analyzed 6 weeks after transplantation by histology and immunohistochemistry. The CRX-mCHERRY+ transplants at this stage were found to be incorporated as large multicellular clusters within the host tissue expressing the characteristic pan-PR-specific marker RCVRN, which also labels the neighboring CRX-mCHERRY-negative host PRs (Fig.2B). Transplants were reproducibly observed in 80% of samples (n=44 transplanted HROs hosts were analyzed out of N=4 separate rounds of donor and host HRO differentiation, on average 11n/N) (Fig.2C). In 86% of the samples with a visible transplant, we found CRX-mCHERRY+ PR cells located in the outer portion of the host retina (Fig.2C), 80% of those with incorporated large multicellular clusters and not as separately integrated dispersed single cells. In the remaining samples (14%), transplants were still ectopically attached to the apical HRO surface (Fig.1E8). To evaluate the proportion of the transplant area incorporated into the hosts, we used a method previously established in animals (14). In 38% of cases with transplant incorporation, 80 to 100% of the transplant area was incorporated into the host retina (n=32, N=4; Fig.2C). In 32%, 20-80% of the transplant area was incorporated. Transplants with only a small proportion of incorporated area (5-20%) were not observed. This supports that as early as six weeks after subretinal-like transplantation, the majority of the transplant has been incorporated into the host.

**Fig.2.**
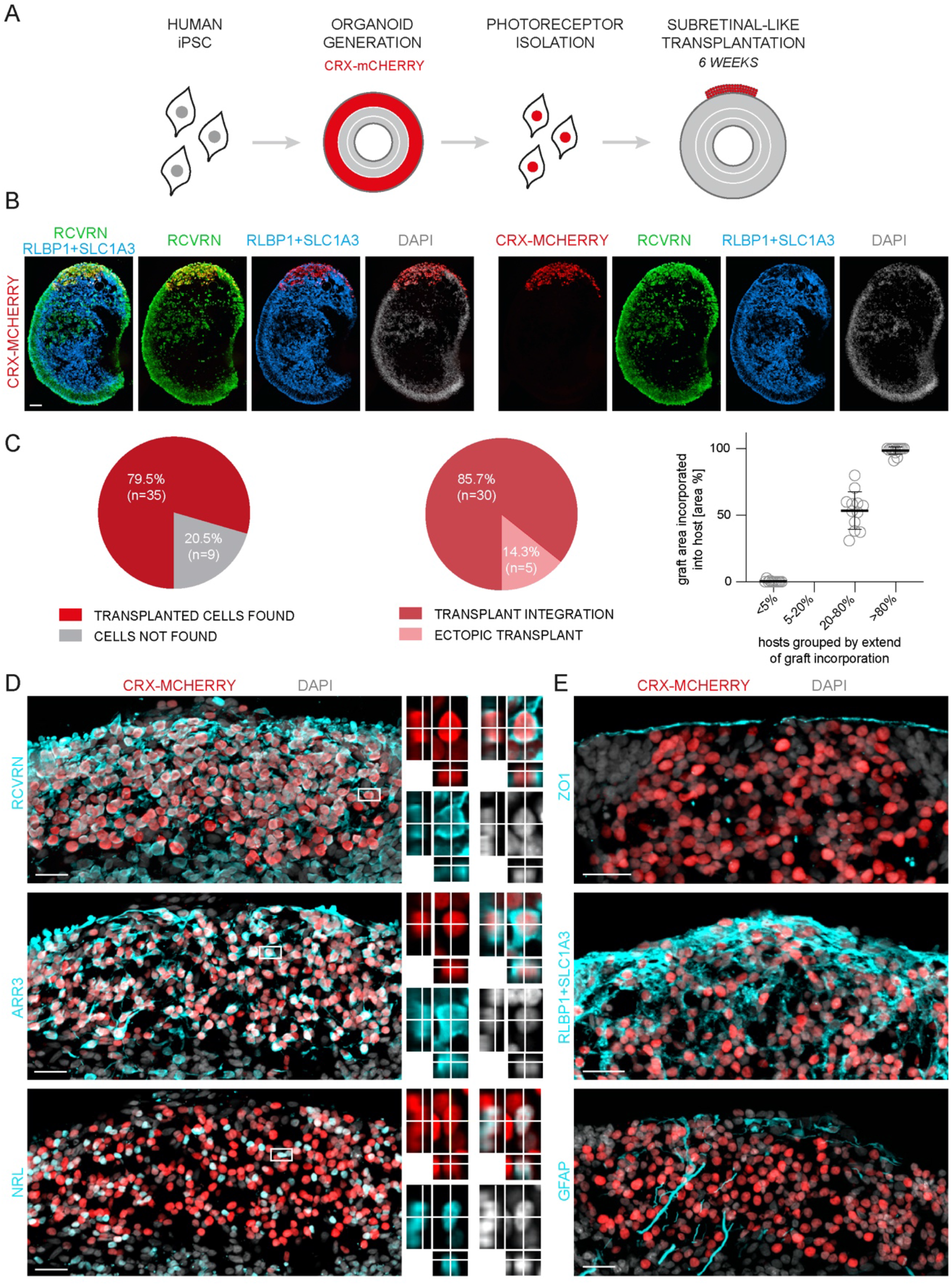
Photoreceptors effectively incorporate six weeks after subretinal-like transplantation into healthy human retinal organoids. (A) Human CRX-mCHERRY+ photoreceptor cells were isolated from retinal organoids and applied for subretinal-like transplantation into healthy human organoid hosts without transgenic reporter. (B) CRX-mCHERRY+ cells reproducibly incorporated as large cell clusters into the host organoid six weeks after transplantation. (C) Transplant PR cells integrated in the majority of samples with a visible transplant (n=44, N=4). In the majority of cases, 20-80% of the transplant area was incorporated into the host (n=32, N=4). Transplants with only a small proportion of incorporation (5-20%) were not observed. (D) Incorporated photoreceptor cells expressed photoreceptor-specific markers (RCVRN); both rod (NRL) and cone (ARR3) photoreceptors were incorporated. Magnifications show stack projections from selected cells to proof threedimensional marker overlap. (E) Cell clusters were infiltrated by Müller glia (RLBP1+SLC1A3) processes, supporting incorporation into and interaction with the host while causing only slight Müller glia reactivity (GFAP). ZO1 cell junction staining indicates partial restoration of the apical epithelial border after transplant incorporation. Scale bars 50*μ*m (B), 25*μ*m (C).

Previous animal studies showed that material transfer of fluorescent reporter from mouse (11–13, 17) but not human (14) PR transplants to host PRs may be misinterpreted as integrated PR transplants. We used a dual reporter approach to assess this: CRX-mCHERRY+ human PRs served as donors, HRO hosts were derived from hiPSCs that ubiquitously express transgenic nuclear enhanced GFP in all cells (NLS-eGFP) (Fig.3A, B), so that eGFP and mCHERRY double-positive nuclei would indicate material transfer. Again, after six weeks transplants were frequently located inside the retinal epithelium of host HROs. CRX-mCHERRY+ and eGFP+ nuclei did not overlap, supporting absence of material transfer, but instead PR integration in human-to-human cell transplantation.

**Fig.3.**
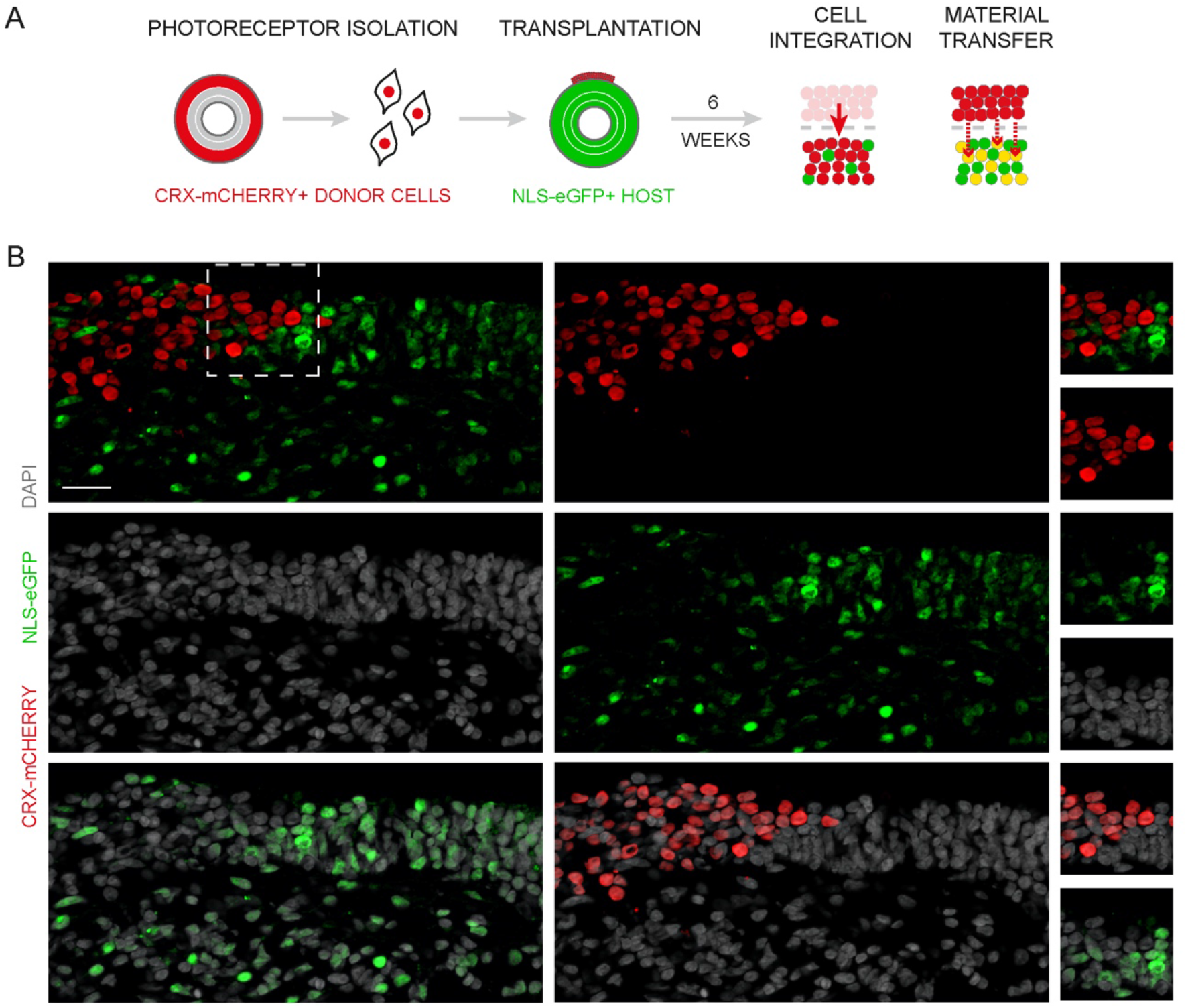
Material transfer between human retinal cells is not evident using a dual nuclear reporter transplantation approach. (A) CRX-mCHERRY+ organoid-derived cells (red) were transplanted into NLS-eGFP+ host organoids (green) and analyzed 6 weeks after transplantation for reporter overlap (yellow), indicative for material transfer between donor and host cells. (B) mCHERRY and eGFP reporters did not overlap, indicating absence of material transfer, but instead PR integration in a human-to-human cell transplantation. Scale bar 50*μ*m.

### Rod and cone photoreceptor transplants exhibit features of structural integration

To investigate PR transplant incorporation in more detail, we performed histological studies. Analysis of PR subtypes by characteristic cell markers showed that both cone (ARR3+) and rod (NRL+) PRs become incorporated into the host retina (Fig.2D); both appeared at comparable numbers and evenly distributed within the integrated clusters. To determine if transplanted PRs are structurally incorporated into the host tissue, and not just closely attached to the organoid surface, we studied them in conjunction with the host MG. Combined RLBP1 and SLC1A3 staining extensively labels the cytoplasm throughout MG in control HROs (Suppl. Fig.S1), showing the typical embedment of all other retinal cells with MG processes. Analysis of transplanted HROs by these MG markers indicate that grafts are infiltrated by numerous cell processes of host MG (Fig.2E). While in the healthy mature retina PR nuclei are tightly packed, stacked and ordered within the outer nuclear layer, incorporated PR transplants and host MG processes frequently still seem unordered with variable distances between PR nuclei and MG process widths. To explore if transplants induced in the host HROs an activation of MG, commonly associated with cellular changes in retinal pathologies, we studied host MG and donor PR cell distributions and immunostained for glial fibrillary acidic protein (GFAP). In injured or degenerating retina, GFAP is a hallmark of MG response to the pathology (42, 43). We rarely observed CRX-mCHERRY-negative host cell nuclei (DAPI) within the clusters of CRX-mCHERRY+ PR transplants (Fig.2D, E), supporting absence of extensive cell delamination. However, we occasionally observed some GFAP filaments in the transplanted PR clusters (Fig.2E), albeit not extensively, and not in retinal epithelial areas next to transplants or in control HROs (not shown), indicating some response of the host HRO to the transplant. Notably, some GFAP filaments were radially oriented indicative of a radial host MG structure within the transplanted PR clusters. Since cell contacts between the apical tip of radial MG processes with the apical portion of neighboring PRs form together the outer limiting membrane (OLM) of the retina, our data posed the question whether transplanted cells might integrate and contribute to this structure.

To assess whether transplanted cells are located below and possibly become part of the host OLM, we analyzed ZO1 on immunostained HRO sections. ZO1 presented as a single band above the most apical row of cell nuclei in control HRO (Suppl. Fig.S1) and above PR transplants (Fig.2E), indicative of an OLM-like structure as the HRO epithelial border. Ultrastructural studies further support this: To visualize the cell morphology of individual transplanted PRs, a subset of cones among the CRX-mCHERRY+ donor population was virally labeled with a cytoplasmic GFP reporter prior transplantation (Fig.4A). CRX-mCHERRY HROs were transduced with recombinant adeno-associated virus (AAV) vectors with an AAV2.GL capsid (39, 44) carrying a single-strand AAV genome with a gene expression cassette coding for the enhanced green fluorescent protein (eGFP) under control of the human cone arrestin (ARR3) promoter (AAV2.GL-ARR3-eGFP). One week after transduction, a subset of CRX-mCHERRY+ cells presented a cytoplasmic fluorescent eGFP signal (Fig.4B1-2). Virally labeled HROs were acutely dissociated (Fig.4B3), enriched for CRX-mCHERRY+ cells by FACS (Suppl. Fig.S2B), and subretinally-like transplanted. Host HROs were analyzed six weeks after transplantation. About 15% of donor cells were eGFP+ prior to transplantation (Suppl. Fig.S2B), and a large number is found within the incorporated transplants (Fig.4B4-5). Mature PR in the human retina and HROs have a characteristic polarized structure (Fig.4C), with an outer and inner segment above the OLM, adherens junctions between inner segments and MG processes, their soma and nuclei in the outer nuclear layer, and a basal process forming a synapse with bipolar interneurons in the outer plexiform layer. Both histological (Fig.4D) and ultrastructural (Fig.4E) analysis by electron microscopy revealed that some of the eGFP-labeled PR transplants formed mitochondria-rich structures apically of the OLM (Fig.4D1, E1a), revealing presumptive PR inner segments. Latter also form adherens junctions with neighboring eGFP-negative cells (Fig.4E1b), further supporting donor-host cell interactions. Immunostaining showed some basal-process like PR structures, which at their ends overlap with RIBEYE immunostaining, a ribbon synapse-characteristic marker, supporting formation of ribbon synapses (Fig.4D2). Electron microscopy confirmed characteristic structures of synaptic ribbons surrounded by synaptic vesicles in eGFP+ PR transplants (Fig.4E2). Together, PR grafts incorporate as cell clusters within the retinal epithelium of host HROs and survive six weeks after transplantation into host HROs, become intermingled with host MG processes, and have the ability to develop some PR characteristic cell morphological features, including formation of donor-host cell contacts contributing to an OLM-like host HRO epithelial structure.

**Fig.4.**
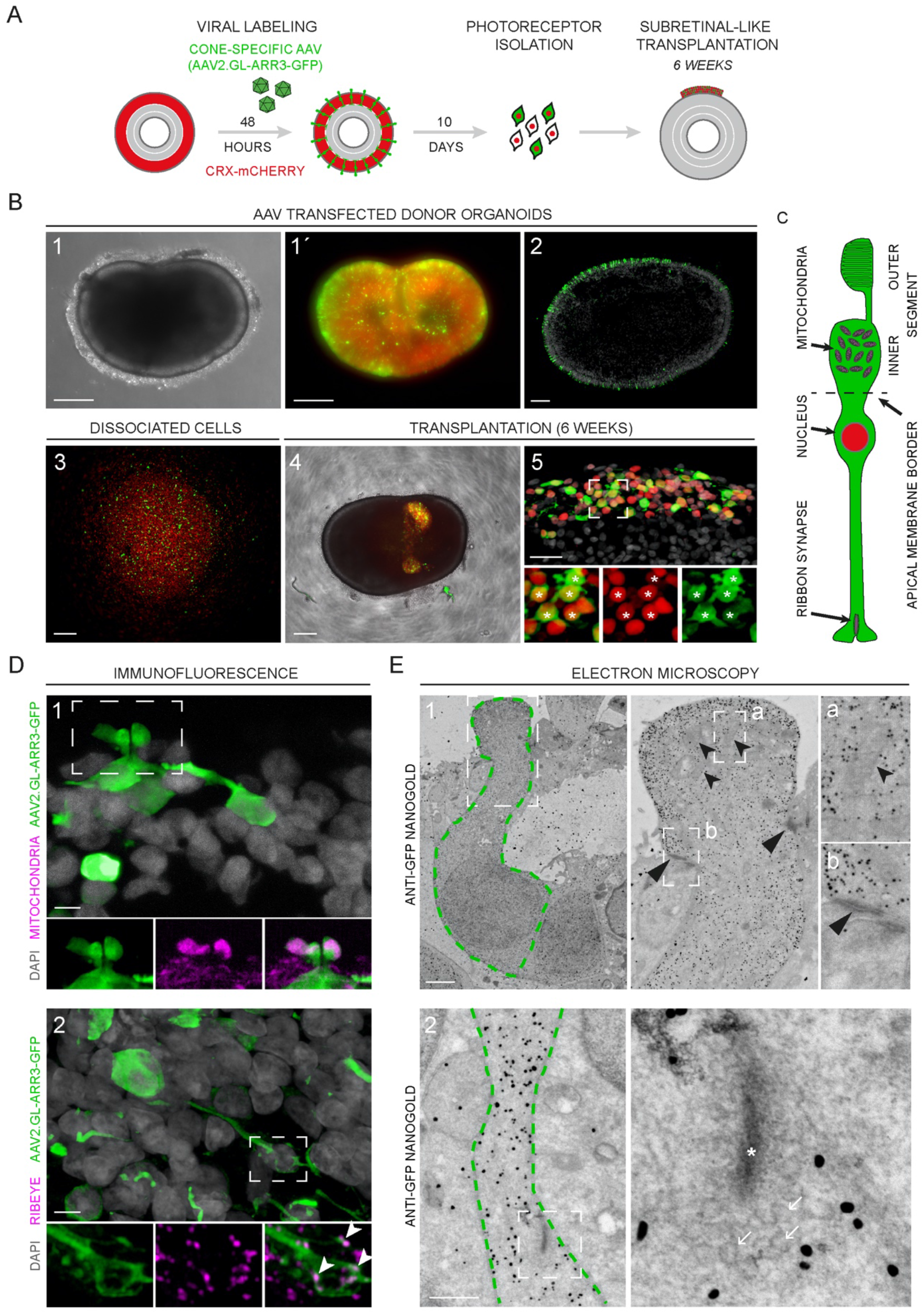
Transplanted human photoreceptor cells exhibit features of structural integration into human organoid hosts. (A) Donor organoids were transduced with adeno-associated virus (AAV) vectors to label a subset of donor cells with cytoplasmic GFP, followed by subretinal-like transplantation for six weeks. (B) 1-3: CRX-mCHERRY (red) and AAV-mediated GFP (green) reporters were expressed in donor organoids and single cells after organoid dissociation. 4: Transplanted cell clusters expressing CRX-mCHERRY and GFP were visible in culture six weeks after transplantation. 5: Incorporated GFP cells were also CRX-mCHERRY+ (asterisk). (C) Cell morphology of a human cone photoreceptor with characteristic ribbon synapse and mitochondria-rich inner segments, located outside the apical border of the retinal epithelium. (D) 1: Mitochondria-rich inner segment-like structures were formed by GFP cells. 2: GFP cell processes overlapped with characteristic ribbon synapse marker (RIBEYE). (E) 1: Ultrastructural analysis of immunogold labeled samples supported the formation of inner segment-like structures containing mitochondria (open arrowheads) by GFP cells as well as formation of cell contacts with GFP negative cells (arrowheads), indicating that they participate in the restoration of the apical epithelial border. 2: Synapses with a characteristic ribbon (asterisk) surrounded by synaptic vesicles (arrows) were observed in GFP cells by electron microscopy. Scale bars 100*μ*m (B, 1-2), 250*μ*m (B, 3-4), 25*μ*m (B, 5), 5*μ*m (D), 1*μ*m (E, 1), 0.5*μ*m (E, 2).

## Discussion

Recent advances in preclinical research on retinal cell replacement therapy in animals provided several evidences that support replacement of cell loss by transplantation of photoreceptors (PRs) for restoration of retinal structure and visual function. Thus, it is time to determine how to effectively translate this potential therapy from animals to patients. A key question is, whether findings in animals can be reproduced also in the human retina with organoids being a promising technology for bridging preclinical animal research towards clinical application.

In the present work, we provide evidences supporting the feasibility of a preclinical human transplantation assay: Human PR transplants can effectively and reproducibly integrate into a human retina model system. We show that transplantation can be achieved by two approaches: Intraretinal- and subretinal-like transplantation of rod and cone PRs as a purified cell suspension derived from postmitotic human retinal organoids (HROs) into 3D healthy host HROs. This results in precisely positioned single PR cells, or structural integration of multicellular PR clusters into the host retina, respectively. To validate transplant integration, we present a dual color transgenic nuclear reporter approach to assess material transfer between donor and host PRs. Our data show absence of material transfer supporting transplant integration. We effectively tracked PR transplants: Both en bulk by transgenic nuclear labeling, showing integrated cells in larger clusters or as single PR cells, and at the individual cell level upon combined cytoplasmic sparse viral-labeling. Thereby, we show that some transplanted PRs acquire several characteristic (ultra)structural features of mature PR in vivo: a polarized cell structure with an apical mitochondria-rich inner segment, soma and nucleus in the outer retina, and a basal process with a synapse-like structure. Previous studies showed that PR in HRO develop axon terminals with specialized ribbon synapses (27, 28, 45). While some PRs integrate into the host retinal barrier, the OLM, by forming donor-host cell contacts with neighboring Müller glia (MG), others still appeared as an amorphous mass six weeks after transplantation. Our data suggest that human PRs can reproducibly be incorporated into a healthy human retina model without additional external stimulus. This supports previous studies of PR transplants into animal models, and offers application potentials and hypotheses to possibly facilitate clinical translation. Future studies will be aimed at determining to which extend transplanted cells may restore cellular structure and function in this experimental system in longer-term experiments, whether transplant integration is influenced by specific pathologies, and whether findings can be reproduced in primary patient retina.

So far, it is still unknown if retinal cell transplants can integrate into patient retina in vivo, as observed in animal models. Our data support this possibility: We show that transplanted PRs integrate into and some contribute to host retinal structure. However, to get from transplant integration to finally improved visual function in patients, several steps not yet assessed in HROs might need to be accomplished on the cellular level (Suppl. Fig.S3): First, transplanted PR need to be incorporated into the host. Second, cell characteristic structural and morphological properties need to be formed, which are the basis to form donor-host cell contacts: Adherens cell junctions with MG cells at the apical boundary of the retinal epithelium and synaptic connections to second order bipolar neurons. Third, cells need to mature and acquire functional properties.

Here, we show human PR transplant integration after 6 weeks into healthy HROs, which is faster than previously described for human PR transplantation into degenerating mouse models in vivo. Notably, comparable experiments in healthy animals have not yet been performed. In degenerating mice, human PRs extensively incorporated after 10 weeks onwards but not yet at 3 weeks, and the degree of structural and functional integration increased over time until functional restoration could be observed at 26 weeks (14). Of note, in latter mouse study and in our work the same CRX-mCHERRY hiPSC line and HRO protocol was used to generate donor PRs for transplantation. Still, the cluster integration is unlikely to be a cell line-specific phenomenon since both studies used a second hiPSC line with a similar integration phenotype. Current data indicates that the donor cell species strongly alters the transplantation outcome. Further, although the integration phenotype of human PRs appears comparable between mouse and human hosts, the timing and mechanism of integration might still differ, host/donor age might play a role, and a pathology might slow down or increase integration. The quality of transplant integration that we observed might be comparable to a mid-stage of integration observed in mice (14). Longer-term HRO transplant studies are required to determine to which extend transplants integrate, and comparative studies might reveal model differences.

HROs are a reduced model system of the retina, which can be advantageous but also limiting for cell transplantation modeling. For example, HROs at day 200 are postmitotic, and contain all major retinal cell types (39). This facilitates studies on the direct primary interaction of transplants with the host retina, which might allow to reveal basic principles of and minimal requirements for cell integration due to its reduced complexity. While other parts, like the retinal pigment epithelium (RPE) opposed to the neural retinal epithelium, microglia or vasculature, are missing in HROs (46), the currently reduced system allows us to determine if they are required for complete transplant integration (Suppl. Fig.S3). Possibly, future implementation of additional components might facilitate functional maturation and wiring of host HROs and cell transplants. Further, our study shows that transplanted PRs may have some pre-synaptic components, thus, it will be interesting to determine if these can form synaptic connections to second order bipolar neurons, specifically, even in a healthy retina setting. Retinal wiring has not yet been extensively studied in HROs. Specifically, retinal ganglion cells become rather rare in postmitotic HROs (28, 29, 31, 47), and are obviously not connected to higher brain centers, but could be in the future (48, 49), which ultimately might be useful to determine if PR transplants can restore function. Notably, HROs at day 200 might have a higher competence and plasticity for cell transplant integration compared to more mature and adult retina. Previous studies showed that human donor PRs derived from day 200 HROs effectively integrate into adult mouse retina in vivo, whereas day 250 old PRs did not (14). Thus, younger HRO hosts might also have properties designating them more permissive for PR transplant integration, which could change upon further maturation and reverse with aging and pathology (50). HROs generated out of patient-derived hiPSCs would allow to apply inherited pathology models as hosts (46, 51). If so, our assay might provide access to the underlying mechanisms facilitating and limiting transplant success at the donor and host level at different ages.

Our data show that in the large majority of cases subretinal-like PR transplants reproducibly integrate in large cell clusters. Only in some cases, PRs integrate sparsely as single cells. In about 25% of cases, we cannot find any transplant in or attached to host HROs, which might be a technical limitation. For example, donor-host contact might become impaired or the transplant might be lost during cell culture medium changes. Alternatively, some HROs might not be permissive for transplant integration. Interestingly, also in animal studies some transplanted PRs remain in the subretinal space and do not contact or integrate into the host retina (14). While HROs under control condition do not show extensive cell death and glial pathologic responses, like reactive gliosis (42, 43), we have not systematically assessed this. Glial pathologic processes, specifically scar formation and neural remodeling at advanced disease stages, is anticipated to present a major barrier to cell replacement strategies (52–55). However, this is not solved, and it is not known yet if transient cell stress or low-level damage, possibly involving responses of MG to the transplant, might prevent its integration.

Previous studies in mice and our data in HROs show that cell processes of host MG extend into and form contact with PR transplants, supporting the notion that MG provide a permissive and supportive (56) rather than repressive environment. PR transplants in HRO at the timepoint studied are mostly unordered, and previous studies in mice showed a comparable histological phenotype at an early stage of transplant integration (14). However, such phenotype is also reminiscent of some retinal pathologies. For example, most pathologies are associated with a response of the MG, called reactive gliosis, which may have beneficial and detrimental functions (56). Gliosis may involve cell hypertrophy, hyperplasia, and displacement. Further, MG may form scars. In degenerating retina, glial fibrillary acidic protein (GFAP) becomes strongly upregulated in MG, a hallmark of reactive gliosis. In the present work, GFAP was observed within the integrated transplants. In previous animal studies, GFAP was also expressed in the PR transplant area (14). Thus, it will be interesting to determine if some level or functions of gliosis might support or limit transplant integration. Further, previous studies in mice showed different phenotypes of PR transplant interaction with the host retina, in the best case with extensive PR incorporation, but also partly integration, and formation of rosette-like epithelial structures reminiscent of outer retinal tubulations frequently observed in retinal pathologies (14). For example, extensive pathologic changes are associated with retinal remodeling (57), like displacement of neurons from the different retinal layers. Notably, in HROs we did not yet observe evidences indicating that MG might form a scar barrier, rosette-like structures, or displacement of entire host cells into the transplants. Instead, MG cell processes intermingled throughout the transplants support donor-host interactions and structural integration.

Comparative and complementary studies of animal models and HROs might further advance cell transplant therapy development. For example, previous studies have revealed species-specific differences of transplant integration comparing mouse and human donor cells. This might also apply for hosts, as the mechanisms regulating transplant integration are still unclear. Even though the healthy mouse and human retina has an intact apical epithelial boundary formed by cell junctions, these might need to be opened up in order to incorporate the transplants. Whether the donor cells induce a loss, reduction, or merge of epithelial boundary components is still unknown. Some type of studies to decipher such question could be performed easier in an ex vivo or in combination with an in vivo system. While primary retina derived from animals rapidly degenerates and develops reactive gliosis in cell culture (43, 58), we did not observe reactive gliosis in control HROs. Thus, HROs might facilitate longer-term live imaging studies recording the dynamic of cell integration processes across several stages (Suppl. Fig.S3). This might also provide insight if donor cells invade the host, or if host cells, like MG, might actively extend processes and take up donor cells. As another example, the subretinal region in the eye is immune privileged, but transplant rejection requiring immune suppression is a limiting factor in animals (14, 59–61). Studies in the HRO system might advance our understanding about the donor-host compatibility in a human setting. The environment of HROs can be tightly controlled, whereas in vivo models have the advantage to study the whole organism including blood and immune system, aspects that are usually missing in organoid models (46). HROs provide higher operational freedom for multiple timed complex manipulations, like extrinsic applications of modulators to improve transplant integration at different stages. In contrast, the mouse eye is a rather small system, and multiple injections pose the risk of inflicting damage and infection. However, studies in mice are advantageous for in vivo functional and behavioral tests to study recovery of visual function (15, 47), while HROs are theoretically available at unlimited numbers and potentially more accessible for invasive functional studies, like patch-clamp electrophysiology, and longer-term functional live imaging, e.g. by transgenic calcium reporters.

The presented human transplantation assay does not require primary patient material which is difficult to acquire with short postmortem times and high quality, and especially from healthy donors, or with pathologic stages optimal for therapies. While our finding that human PR transplants integrate into HROs is encouraging and in line with previous animal studies, reproduction of key data in primary human retina ex vivo will be the next step towards validation of key findings and thereby of the HRO transplantation assay. For example, primary retina might enable studies of PR integration in adult retinal tissue since ageing is still a limiting factor of HRO technology (62). However, even though HROs are a reduced system, these might even more closely reproduce a healthy in vivo setting ex vivo compared to culture of primary retina, which spontaneously degenerate ex vivo (43, 62). With the availability of organoids from different organisms (63, 64), comparison of other species-specific differences might become possible, for example, to determine large animal models optimal for in vivo studies, which might be useful to optimize surgical methods for clinical translation (65–67). Further, the herein established HRO transplantation assay might advance several remaining major questions in cell replacement therapy development towards clinical translation. For example, depending on the pathologic conditions, defined cell products will be required for cone and/or rod PR transplants that do not become rejected. Another example, to define types and stages of pathologies that will be optimal targets for effective transplant integration, as well as visual restoration, and possibly prevention of visual function loss. Thus, in future studies HROs with defined pathologies could be utilized to further develop and optimize cell transplant integration. Thus, the HRO transplantation assay might be useful to facilitate cell replacement therapy development for age-related macular degeneration, inherited retinal dystrophies, glaucoma, and other pathologies.

In summary, we provide a proof of concept for the integration of transplanted human rod and cone PRs into a human retina ex vivo model, which might provide a preclinical experimental human cell transplantation system facilitating retinal cell replacement therapy development, translation, and clinical optimization. The presented organoid-based preclinical human cell transplantation system might be of interest not only for retina research but for other cell therapeutic approaches, and the presented method could be directly applied for some other types of tissue specific organoids.

## Acknowledgment

We thank the DZNE light microscopy facility as well as the core facilities of the CRTD and CMCB technology platform at TU Dresden for excellent support: CMCB light microscopy facility, stem cell engineering facility, flow cytometry facility and electron microscopy facility. We thank the whole MOKALAB, especially the organoid team, and Anne Wagner, for excellent support. We thank Marius Ader (CRTD, TU Dresden) for helpful comments on the manuscript, and Sylvia Gasparini and Karen Tessmer for sharing technical assistance and expertise. We thank Botond Roska and George Church for providing plasmids for the generation of the CRTD1-NLS-eGFP hiPSC line.

## Authors contributions

Conceptualization, F.W. and M.K; Methodology, F.W., M.Z., V.B. and M.K.; Formal Analysis, F.W. and T.K.; Investigation, F.W., R.C., M.Z., V.B., T.K, and R.N.; Resources, S.M., K.N. and O.G.; Writing – Original Draft, F.W. and M.K.; Visualization, F.W. and T.K.; Supervision, F.W. and M.K.; Project Administration, M.K.; Funding Acquisition, M.K.

## Funding

Supported by the Funding Programs for DZNE Helmholtz (M.K.); TU Dresden CRTD (M.K.); Deutsche Forschungsgemeinschaft (DFG, German Research Foundation): SPP2127 KA2794/5-1, KA2794/5-2 (M.K.); HGF ExNet-007 (M.K.); Bundesministerium für Bildung und Forschung (BMBF): ReSight (01EK1613A); ERA-NET Neuron ReDiMoAMD (01EW2106); EFRE (EM Facility, T.K.).

## Declaration of interests

OG is inventor on patents on hiPSC retinal differentiation (patent nos. EP2989200/WO2018149985) and on the use of hiPSC retinal derivatives to treat retinal degeneration (patent no. WO2018055131), licensed to Gamut Cell Tx. All other authors declare no competing interests.

## Methods

### Cell lines

All procedures involving human induced pluripotent stem cells (hiPSC) were performed in accordance with the ethical standards of the institutional and/or national research committee, as well as with the 1964 Helsinki declaration and its later amendments, and approved by the ethical committee at the University of Dresden.

For differentiation of human retinal organoids, the following hiPSC lines were used in this study: 5A (68), CRX-mCHERRY (21) and the newly generated CRTD1-NLS-EGFP line. All hiPSC lines were maintained in mTeSR1 (Stem Cell technologies) on Matrigel-coated culture dishes at 37°C and 5% CO_2_ and passaged using ReLeSR (Stem Cell technologies). CRX-mCHERRY hiPSCs were grown on vitronectin (VTN-N; Thermo Scientific).

### Generation of nuclear CRTD1-NLS-EGFP reporter hiPSC line

The NLS-EGFP cassette was PCR-amplified using KAPA HiFi HotStart ReadyMix (Roche) using the plasmid pAAV2-CAG-ChR2d-2A-NLS-EGFP (kind gift of Botond Roska) as a template. The primers were decorated with 5’ BamHI and 3’ BsrGI linkers. Following primer were used: For-NLS-ggatccggatccgccaccatgactgctccaaagaagaag and Rev-NLS: ttacttgtacagctcgtccatgccgagagtgatcccggc. The piggybac backbone PB-eF1a-EGFP-T2A-Puro (generated in the Busskamp lab, parental vector Addgene plasmid ID #63800, kind gift of George Church) was linearized using BamHI and BsrGI restriction enzymes. Subsequently, the BamHI/BsrGI digested PCR product was ligated to the linearized backbone using the DNA Ligation Kit - Mighty Mix (TaKaRa) resulting pPB-ef1a-NLS-EGFP-T2A-puro.

The nucleofection of the pPB-ef1a-NLS-EGFP-T2A-puro cassette into CRTD1 hiPSCs (hPSCreg: CRTDi004-A) was performed as previously described (69). Briefly, 10 *μ*g of pPB-ef1a-NLS-EGFP-T2A-puro and 2 *μ*g of Super PiggyBac Transposase Expression Vector (Biocat) were electroporated into CRTD1 cells with the Lonza 4D X-unit, pulse CB-150 using the P3 Primary Cell 4D-Nucleofector Kit L (V4XP-3024, Lonza). CRTD1 -NLS-EGFP cells were selected using puromycin.

For Fluorescent cell sorting, CRTD1-NLS-EGFP hiPSCs were dissociated by TrypLE Express (ThermoFisher Scientific), resuspended in cold mTeSR Plus medium with CloneR and immediately sorted using a FACS Aria III (BD). EGFP positive cells with a moderate GFP signal were seeded to Matrigel-coated 96-wells in mTeSR Plus medium supplemented for 3 days with CloneR (both StemCell Technologies). After 3 days, half of the medium volume of mTeSR Plus with CloneR was freshly added on top without removing medium from wells. After 5 days medium was changed completely once before switching to mTeSR1 medium on day 7 for regular culture. Single clones were expanded by ReLeSR passaging. From the second passage after sorting on 2 *μ*g/ml puromycin were added for 24h after each split to avoid silencing of the transgenes. A clone with a homozygous and moderate GFP expression was selected for characterization and a master cell bank was cryopreserved.

### Human retinal organoidogenesis

Human retinal organoids (HROs) were differentiated from previously published hiPSC lines (see section: Cell lines): 5A and CRX-mCHERRY using our previously published protocol (39). Briefly, hiPSCs were passaged into small clumps and embedded in Matrigel for solidification. The gel was dispersed into small clumps and cultured in N2B27 medium (1:1 DMEM/F12:neurobasal A medium, 1% B27+Vitamin A, 0.5% N2, 1% penicillin/streptomycin, 1% GlutaMAX, 0.1mM 2-mercaptoethanol). Neuroepithelial cysts spontaneously formed within a few days. On D5, cysts were plated on Matrigel-coated plates. On D13 adherent cysts were detached using dispase and further cultured in B27 medium (DMEM/F12, 1% B27 without vitamin A, 1% penicillin/streptomycin, 1% GlutaMAX, 1% NEAA, 0.1% amphotericin B), supplemented with 10% FBS from D25 on. Retinal domains were manually isolated between D25 and D30 using surgical tweezers. On D100, medium was changed to N2+FBS+DMEM/F12 (1% N2, 10% FBS, 1% penicillin/streptomycin, 1% GlutaMAX, 0.1% amphotericin B). Medium was supplemented with EC23 (0.3*μ*M) from D25 to D120, half medium was replaced every 2-3 days. Culture conditions were 37°C and 5% CO_2_ for the whole time period.

### Organoid dissociation and flow cytometry

HROs from the CRX-mCHERRY hiPSC line (D170–D220), unless stated otherwise, were dissociated using the Papain Dissociation System (Worthington Biochemical Corporation, LK003150). Reagents were prepared according to the manufacturer’s instructions, however, DNase I was substituted (Sigma-Aldrich, D5025-150KU). 10–15 organoids were pooled and washed with PBS (37°C, 3x). PBS was replaced with 1ml preheated papain (20 U/ml, 37°C), followed by a 5 min incubation at 37°C and 300 rpm (Eppendorf Thermomixer comfort). HROs were incubated on an orbital shaker incubator (ES-20/60, Biosan, 37°C, 90rpm, 2 hours). 50*μ*l DNase I was added and allowed to sit for 5 minutes at room temperature. The samples were slowly triturated ~10x with a fire-polished glass pipette until a homogenous suspension was obtained. The cell suspension was added to a wash solution containing 250*μ*l EBSS, 60*μ*l Ovomucoid and 60 *μ*l DNase I. The mixture was then overlaid onto 1ml of Ovomucoid and centrifuged for 6 min. at 600 g at room temperature. After centrifugation, the cells were resuspended in 1-2ml of MACS buffer (2mM EDTA, 0.5% BSA in PBS) and filtered (35*μ*m mesh) into flow cytometry tubes (Falcon, 352235). Afterwards, 20*μ*l/ml of DNase I was added and cells were kept on ice until FAC-sorting. CRX-mCHERRY+ cells were sorted with a BD FACS Aria III cell sorter using a 100*μ*m nozzle. The cell population was enriched for photoreceptors containing the CRX-mCHERRY+ reporter by 651 nm laser excitation. DAPI was applied (1:1000) right before the FAC-sort (405 nm laser excitation) in order to exclude dead cells and membranous debris. A short re-analysis of the sorted cells was done to corroborate sample purity. 20*μ*l/ml of DNase I was added and the tube was placed on ice for transport. The cells were centrifuged (600g, 6 min, 4°C) and resuspended in 1ml of N2+FBS+DMEM/F12 medium. The cell number was determined with an automated cell counter (BIO-RAD, TC20). On average, we isolated 42,000 cells per HRO (n=126, N=3).

### Intraretinal cell transplantation

Injections of FAC-sorted CRX-mCHERRY+ cells were performed on a Zeiss Axiovert 200M microscope with a self-built glass chamber as sample holder that was cooled to 10°C. A left- and right-sided Narishige MMO-4 manipulator was used for manipulation. The left side was used to fix the host HROs with a holding needle in negative pressure using an Eppendorf Cell Tram Air. The holding needle had an outer diameter of 300 *μ*m and was melted down to about 100 *μ*m with a microforge to gently fix the HROs.

With the right side, photoreceptor cells were injected into the retinal epithelium of the host HROs. We used an Eppendorf Zell Tram Vario system to manually collect and inject the FAC-sorted CRX-mCHERRY+ donor cells. The injection needle had an inner diameter of 15 *μ*m and an angle of 45 degrees at the tip. Since both capillaries have angled needle mounts on the manipulator, both capillary ends (approx. 2 mm) were bent to approx. 30 degrees, so that they work parallel to the glass chamber.

Only donor cells that appeared healthy, based on their cell shape, were collected. After sample immobilization with the holding needle, the injection needle was moved into the bright epithelial part (presumptive photoreceptor layer) of the organoid, visible by phase contrast. The capillary was adjusted to penetrate the organoid in a small angle, which was almost parallel to the organoid surface border. 30 CRX-mCHERRY+ cells were carefully injected. The injection capillary was slowly moved out of the organoid during the injection procedure. Due to the small injection angle, this allowed distribution of the single cells along the epithelial length instead of creating one large cell cluster. This approach was performed with 5 HROs in total. Samples were fixed 3 weeks after transplantation.

### Subretinal cell transplantation

35,000 FAC-sorted CRX-mCHERRY+ donor cells were seeded into each well of a 96-well low attachment plate (Lipidure) in 180 *μ*l N2+FBS+DMEM/F12 medium. Cells settled in the center of the well within 2-3 hours. Then, one >180-days old host HRO was gently transferred into each well on top of the donor cells using wide opening pipet tips. Host HROs from 5A line without fluorescent reporter were used unless stated otherwise. To not impair the aggregated donor cells, HROs were allowed to sink down from the pipet tip into the well without pipetting. Host HROs were selected based on a roundish shape without multiple retinal domains to standardize and optimize host-donor interaction. Half medium was changed every 2-3 days using Xplorer multichannel pipettes (Eppendorf) set at lowest speed to not disturb the host-donor interaction. Samples were fixed after 6 weeks.

### Tissue preparation

HROs were fixed in 4% PFA in PBS for 30min at room temperature (RT). For cryosections, HROs went through a graded series of sucrose (10%, 30%, 50%) and were embedded in O.C.T. (Sakura Finetech). Before freezing in O.C.T., HROs were oriented under an inverted fluorescence microscope (Echo Revolve) so that the cell transplant faced the bottom of the cryomold. Serial cryosections at 12*μ*m thickness were prepared using a CryoStar NX70 cryostat (Thermo Scientific). Samples were oriented so that cross-sections of the transplant region were generated. Tissue slices were mounted on Superfrost Plus slides (Thermo Scientific) and stored at −80°C.

### Immunohistochemistry

Sections were warmed to room temperature and rehydrated in 1X PBS for 15min. If necessary, an antigen retrieval step was performed (10mM sodium citrate, pH 6.0, 30min at 70°C). Blocking solution (0.5% BSA and 0.3% Triton-X-100 in PBS) was applied for 1h at room temperature followed by primary antibody incubation for 48h at 4°C. Primary antibodies used in this study are listed in the key resources table. Washing steps were performed (3x 15min; 1X PBS), followed by incubation with species-specific secondary antibodies conjugated to fluorophores (AF488, Cy3, AF647; Dianova; 1:1000) for 1h at room temperature. If necessary, phalloidin was stained for 30min at room temperature. Cell nuclei were counterstained with DAPI (AppliChem; 1:10,000 in PBS) for 3min at room temperature. After additional washing steps (3x 15min; 1X PBS), slides were coverslipped with Fluoromount-G (Invitrogen) and stored at 4°C.

### Microscopy

Histological samples were imaged on an upright Zeiss microscope with an ApoTome2 slider using Plan Apo 5x/0.16, Plan Apo 10x/0.45 and Plan Apo 20x/0.8 objectives. DAPI, AF488, cy3 and AF647 fluorophores were excited using an HXP light source and the following filter settings: EX BP G365; BS 395; EM BP 445/50 (DAPI), EX BP 470/40; BS 495; EM BP 525/50 (AF488), EX BP 546/12; BS 560; BP 575-640 (Cy3), and EX BP 640/30; BS 660; EM 690/50 (AF647). Z-stacks with 1*μ*m slice distance were acquired that cover the whole epithelium.

For scoring of transplant integration frequency and measurement of incorporated transplant area, whole slides were scanned using an AxioScan.Z1 (Zeiss) with a Plan-Apochromat 10x/0.45 objective, a LED light source Colibri 7 (Zeiss) with respective filters and an Orca Flash 4.0 V2 (Hamamatsu) camera.

An inverted point scanning confocal microscope (LSM 980, Zeiss) with AiryScan 2 detector, Multiplex 4Y scan mode and 40x/1.2 water immersion objective was used for high resolution imaging of integrated AAV-transduced donor cells. Pixel size was 0.052 × 0.052 × 0.180*μ*m. 405nm, 488nm and 639nm lasers were used for excitation of DAPI, AF488 and AF647 fluorophores, respectively.

### AAV production

Recombinant adeno-associated virus (AAV) vectors were produced with AAV2.GL capsids (44) according to previously described procedures (70). For transduction of donor cells, we used AAV2.GL capsids carrying a single-strand AAV genome with a gene expression cassette coding for the enhanced green fluorescent protein (eGFP) under control of the human cone arrestin (ARR3) promoter (71) (AAV2.GL-ARR3-eGFP). AAV vector preparations were stored at −80 °C until use.

### AAV transduction of donor cells

To evaluate inner segment formation and synaptic connectivity of transplanted photoreceptor cells after integration, donor organoids were transduced with AAV2.GL-ARR3-eGFP to label cone photoreceptors. 200-days old CRX-mCHERRY HROs were cultured in 12-well low attachment plates (Nunc; Thermo Scientific) with 6 organoids per well in 1ml N2+FBS+DMEM/F12 medium. HROs were incubated with AAV2.GL-hARR3-eGFP (3.9×10^9 vg / HRO) for 48h. Viral vector-containing medium was removed by 3 full medium changes. After 9 more days, with 50% medium changes every other day, organoids were used for dissociation and FAC-sorting of CRX-mCHERRY+ cells. Subretinal transplantation was performed as described above. Medium was supplemented with 1% amphotericin B before AAV incubation and throughout the whole transplantation experiment.

### Electron microscopy

Organoids were fixed 6 weeks after transplantation of AAV2.GL-AAR3-GFP transduced photoreceptor cells in 4% PFA buffered in 100 mM phosphate buffer. 50 *μ*m thick vibratome sections were prepared using a Leica VT1200 S. Sections with transplanted photoreceptors were selected using GFP-fluorescence. The vibratome sections were subjected to preembedding immunogold labeling using Nanogold and silver enhancement (72, 73). In brief, the samples were blocked and permeabilized in 20% normal goat serum (NGS) / PBS / 0.1% Saponin for 2hrs at room temperature followed by incubation in primary antibody (rabbit anti-GFP, Torrey Pines, TP401, 1:100) in 20% NGS/0.05% Saponin for 2 days at room temperature. After washes in 20% NGS/0.05% Saponin, the samples were incubated with goat-anti-rabbit Nanogold (Fab-fragments, Nanoprobes, 1:50) in 20% NGS/0.05% Saponin overnight at room temperature. After final washes in 20% NGS/0.05% Saponin, the samples were postfixed in 1% glutaraldehyde in PBS, washed in water, silver enhanced using the SE-Kit (Aurion, 1h incubation time), followed by washes in water, postfixation in 1% osmium tetraoxide/water (1h on ice), washes in water, en bloc contrasting with 1% uranyl acetate (1h, on ice), washes in water, and dehydration in a graded series of ethanol (30%, 50%, 70%, 90%, 96% ethanol/water, 3x 100% ethanol on molecular sieve). The samples were infiltrated in mixtures of the Epon substitute EMbed 812 with Ethanol (1+2, 1+1, 2+1, 2x pure epon), flat embedded on the surface of an empty epon dummy block, and cured overnight at 65°C. The region with the transplanted photoreceptors was identified by correlating the fluorescence images of the freshly cut vibratome sections with the finally cured samples, and the target region was trimmed for ultrathin sectioning. Ultrathin sections (70 nm) were prepared using a diamond knife and the Leica UC6 ultramicrotome (Leica Microsystems, Wetzlar), mounted on formvar coated slot grids, and contrasted with 4% uranyl acetate/water for TEM. Sections were imaged with a Jeol JEM1400Plus transmission electron microscope running at 80kV acceleration voltage.

### Evaluation of material transfer

To evaluate material transfer, a dual reporter approach was pursued. CRX-mCHERRY+ donor cells, acutely isolated from HROs, were enriched by FACS and subretinally transplanted into CRTD1-NLS-EGFP host organoids as described above. Experiments were stopped 6 weeks after transplantation.

### Image processing

Graphs were prepared using GraphPad Prism 9 software. Images were optimized by making minor changes to contrast and hue in Adobe Photoshop. Schemes and figures were prepared using Adobe Illustrator software.

### Scoring of transplant integration

In total, n=44 HRO transplantation hosts were analyzed out of N=4 separate rounds of donor and host HRO differentiation (9-15n/N). Samples were analyzed 6 weeks after transplantation. To quantify the frequency of photoreceptor cell integration into host organoids, Axioscan images of serial HRO cryosections were analyzed 6 weeks after transplantation. An integrative phenotype of the CRX-mCHERRY+ donor cell cluster was assigned when CRX-mCHERRY+ nuclei were found below the apical border of the host organoid, using Müller glia counterstaining (SLC1A3+RLBP1) as a proxy.

### Incorporated graft area analysis

HRO serial sections were prepared and every 9th section out of about 80 was analyzed. To evaluate the degree of transplant incorporation into the host HRO, Axioscan images of histological sections were evaluated 6 weeks after transplantation (n=32 HROs out of N=4). The methodological approach was performed and groups were assigned as described before (14). In brief, a line was drawn at the host apical border judged on Müller glia counterstaining (SLC1A3+RLBP1), and the transplanted cell area was measured above (not incorporated) and below (incorporated) of this line.

### Statistical analysis

Sample sizes were chosen based on our own preliminary studies. HROs were randomly assigned to the experiment. For the purpose of reproducibility, HROs with several retinal domains, and hence a shamrock-like shape instead of an even roundish or elliptic shape, were not selected as hosts HROs for technical reasons explained above. Data are plotted as mean and standard deviation. Means were calculated from total HRO numbers (n) that were derived from N number of separate rounds of host and donor HRO differentiations. All graphs were prepared using GraphPad Prism 9 software.

---

**Suppl. Fig.S1.**
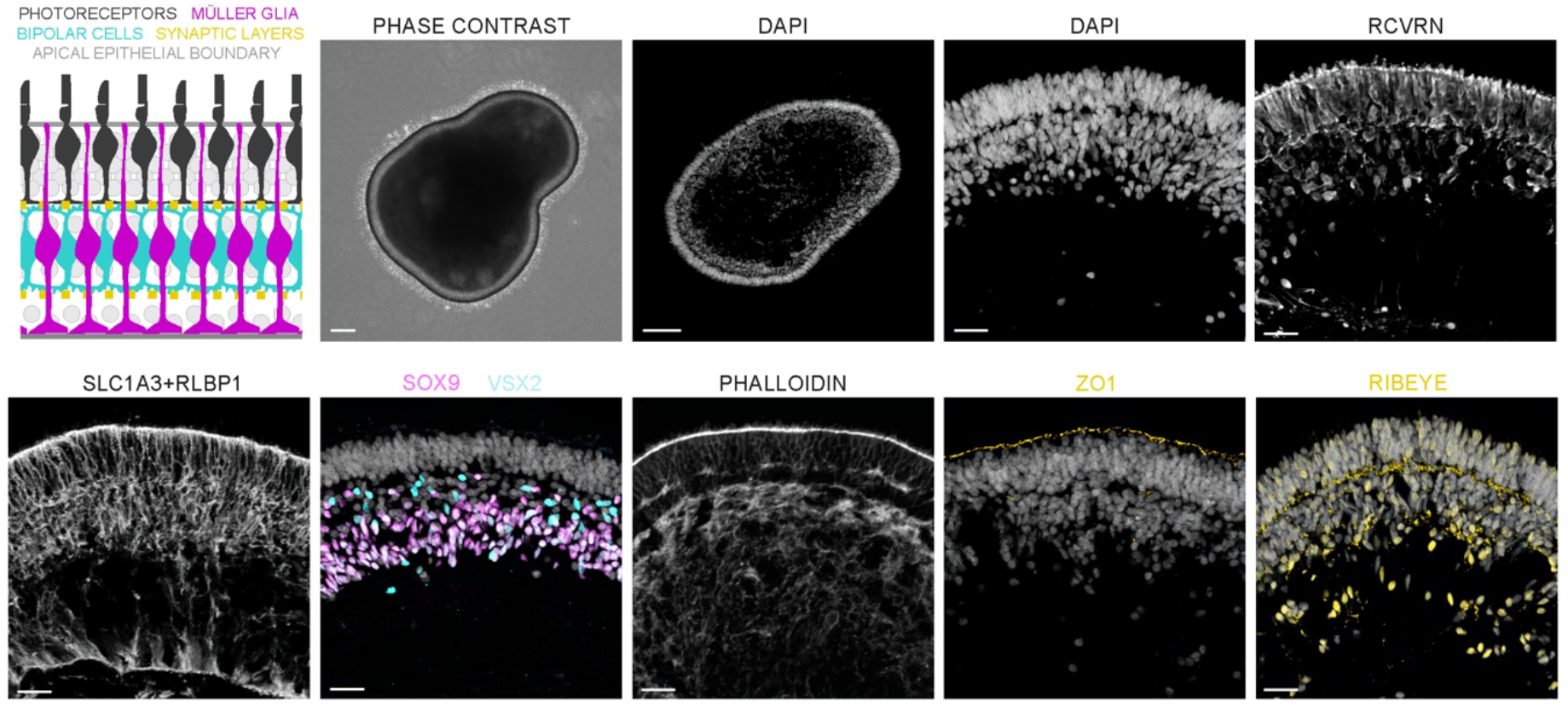
200-days old human retinal organoids (HROs) recapitulate epithelial structure and cell type composition of the in vivo retina. HROs exhibit epithelia with multiple nuclear layers (DAPI) and an intact apical epithelial boundary (PHALLOIDIN, ZO1) formed by Müller glia cell processes (SLC1A3+RLBP1). Photoreceptor cells (RCVRN) are located in the outer retina, Müller glia (SOX9) and bipolar (VSX2+SOX9-) nuclei are part of the inner retina. Inner and outer nuclear layers are separated by a synaptic layer (RIBEYE). Scale bars 150*μ*m (phase contrast), 100*μ*m (DAPI low magnification), 25*μ*m.

**Suppl. Fig.S2.**
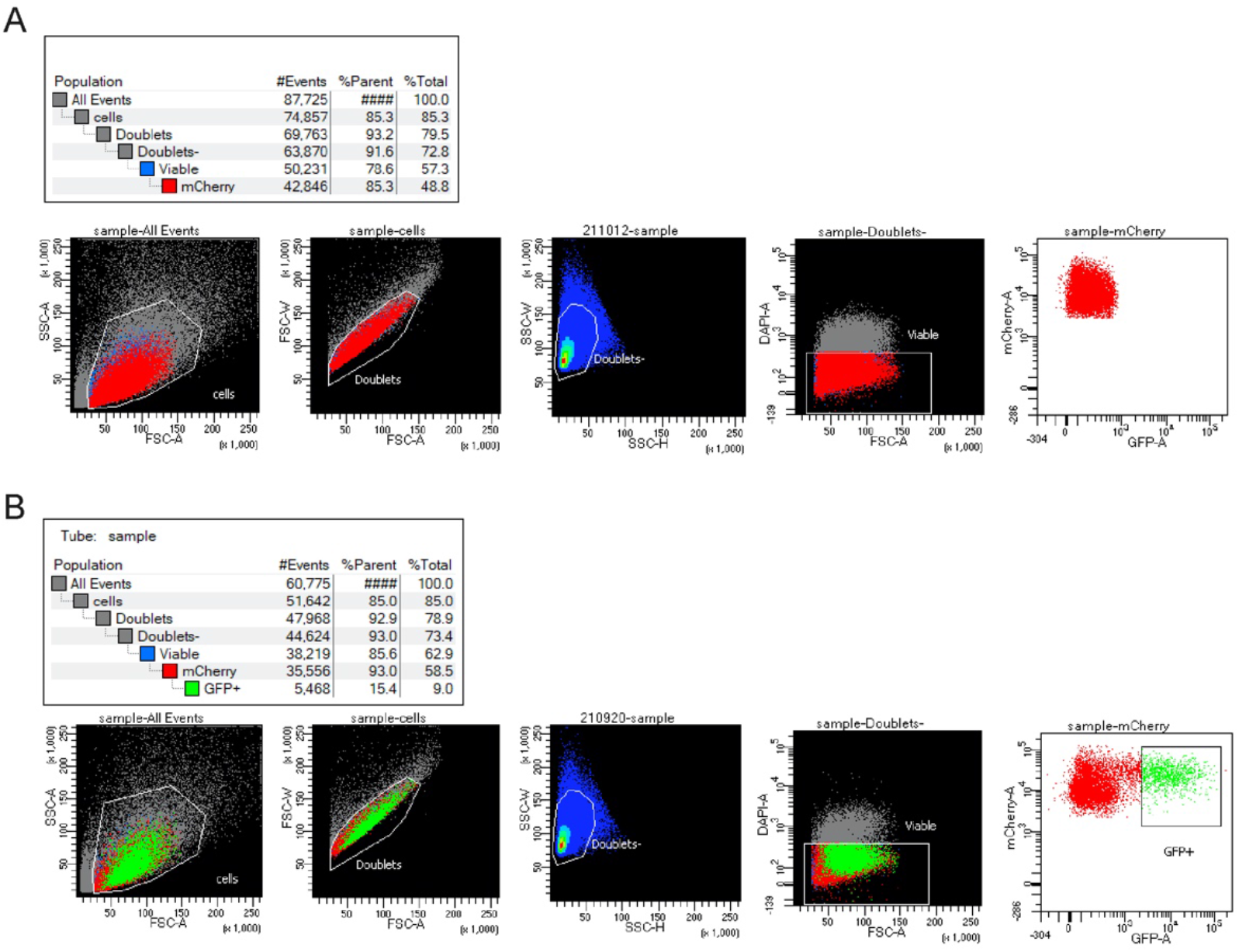
Gating strategy to enrich donor photoreceptor cells by fluorescence-activated cell sorting (FACS) prior transplantation. CRX-mCHERRY+ photoreceptor cells were enriched from dissociated human organoids. Dead cells, debris and cell doublets were excluded during the sort as shown in forward and side scatter (FSC, SSC) plots. (A) Enrichment of CRX-mCHERRY+ cell population from donor organoids. (B) Enrichment of CRX-mCHERRY+ cell population from donor organoids that were in addition virally transduced prior dissociation (using AAV2.GL-ARR3-eGFP) to label a subpopulation of CRX-mCHERRY+ cone photoreceptor cells with cytoplasmic eGFP.

**Suppl. Fig.S3.**
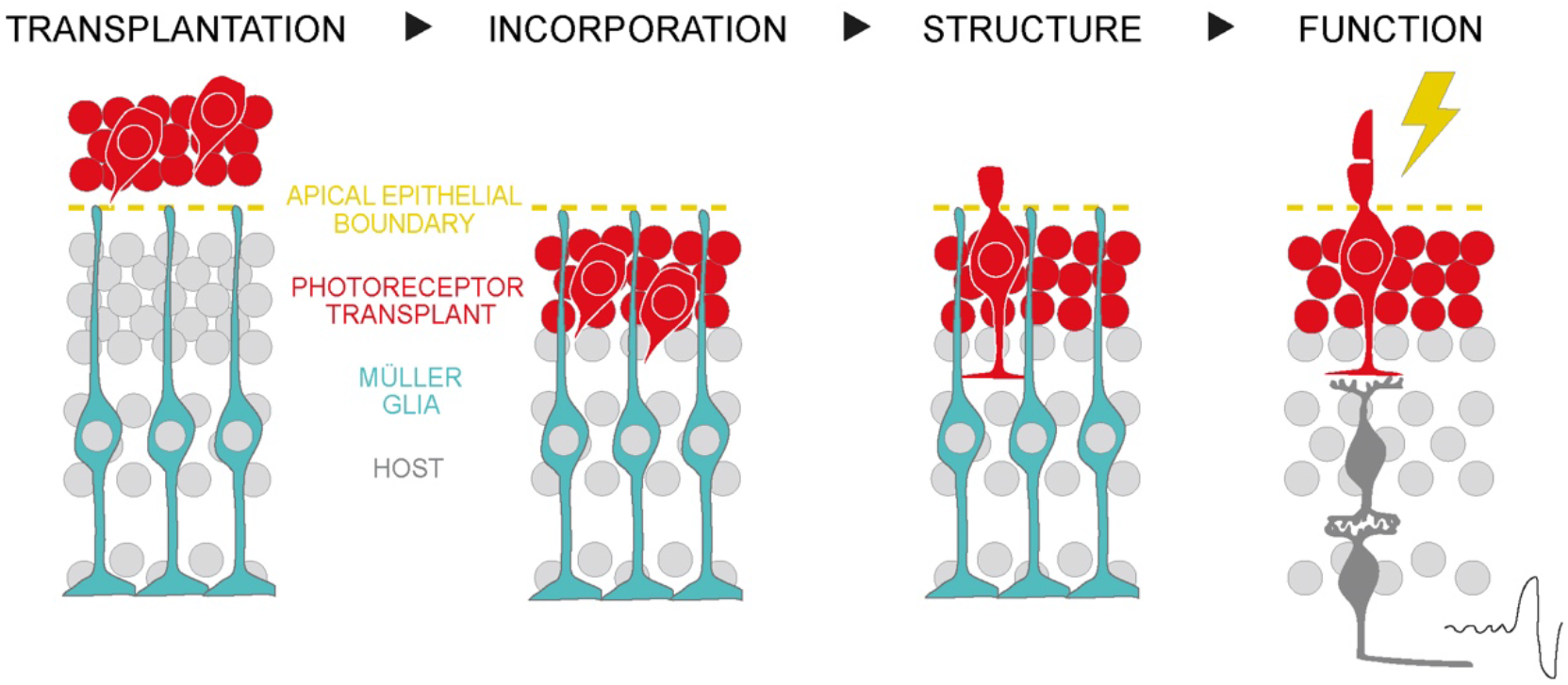
Working model: Stages from photoreceptor cell integration to restoration of visual function. Transplanted photoreceptor cells (red) first need to be incorporated into the host (gray nuclei) by crossing the apical epithelial boundary (yellow line). Second, structural and morphological properties need to be reformed, including interaction with host Müller glia cells (blue). Third, functional integration is required by formation of synaptic connections with host neurons and acquisition of functional properties to transmit light-induced signals and ultimately restore vision.

**Suppl. Movie 1. Collection of photoreceptor cells for intraretinal transplantation.** An injection needle with 15*μ*m inner diameter was used to manually collect single photoreceptor cells derived from dissociated human retinal organoids.

**Suppl. Movie 2. Intraretinal transplantation of photoreceptor cells.** Collected photoreceptor cells were injected into the bright epithelial part of immobilized human organoid hosts.

